# Mechanistic Insights into the Effects of Key Mutations on SARS-CoV-2 RBD-ACE2 Binding

**DOI:** 10.1101/2021.09.20.461041

**Authors:** Abhishek Aggarwal, Supriyo Naskar, Nikhil Maroli, Biswajit Gorai, Narendra M. Dixit, Prabal K. Maiti

## Abstract

Some recent SARS-CoV-2 variants appear to have increased transmissibility than the original strain. An underlying mechanism could be the improved ability of the variants to bind receptors on target cells and infect them. In this study, we provide atomic-level insight into the binding of the receptor binding domain (RBD) of the wild-type SARS-CoV-2 spike protein and its single (N501Y), double (E484Q, L452R) and triple (N501Y, E484Q, L452R) mutated variants to the human ACE2 receptor. Using extensive all-atom molecular dynamics simulations and advanced free energy calculations, we estimate the associated binding affinities and binding hotspots. We observe significant secondary structural changes in the RBD of the mutants, which lead to different binding affinities. We find higher binding affinities of the double (E484Q, L452R) and triple (N501Y, E484Q, L452R) mutated variants than the wild type and the N501Y variant, which could contribute to the higher transmissibility of recent variants containing these mutations.

## Section 1. Introduction

The COVID-19 pandemic has caused more than 4.5 million deaths so far. The severe acute respiratory syndrome coronavirus 2 (SARS-CoV-2), the causative agent of COVID-19, attaches to target host cells with its spike protein, called S, by binding to the angiotensin-converting enzyme 2 (ACE2) receptor, mainly expressed in the lungs^1–4^, and causes the infection. The binding of S to ACE2 triggers conformational changes in S from its metastable prefusion state to a stable post-fusion state^5^. The spike protein is divided into two subunits, S1 and S2, in the prefusion stage. The receptor binding domain (RBD) in S1 is responsible for the binding with ACE2, whereas S2 mediates the subsequent fusion of viral and cell membranes, allowing viral entry^5^. Characterizing the molecular details of the interaction between the RBD and ACE2 plays an important role in understanding the process of SARS-CoV-2 infection of target cells. This understanding may also help explain the improved transmissibility that is seen with SARS-CoV-2 variants that carry mutations potentially affecting RBD-ACE2 binding.

SARS-CoV-2 belongs to the betacoronavirus genus, which includes SARS-CoV and the middle-east respiratory syndrome virus (MERS)^6^. SARS-CoV-2 and SARS-CoV have high sequence similarity, of ∼80%. SARS-CoV-2, however, has spread far more than SARS-CoV and that may be due to the higher binding energy of SARS-CoV-2 with ACE2^7,8^. More recently, several variants of SARS-CoV-2 carrying mutations in RBD, and other regions of S have been reported that have heightened transmissibility and can cause infections with different severity among individuals^9,10^. The double mutated variant B.1.617 has already caused a second wave of COVID cases in India^11^. There have been several reports from the Indian sub-continent about the potential infection and death rate or severe complications among the B.1.618 triple mutant variants^12^. The underlying molecular mechanism behind the binding of these variants with human ACE2 receptor is still unknown.

In this study, we estimate the binding energy between the ACE2-RBD of the SARS-CoV-2 wild-type, and of its single (N501Y), double (E484Q, L452R) and triple (E484Q, L452R, and N501Y) mutated variants. We have considered the two mutations E484Q and L452R seen in the variant B.1.617 responsible for the second wave in India^13^. We considered these mutations along with B.1.1.7 N501Y variant, one of the early mutations argued to have caused increased transmissibility^14^. We calculate the binding energy differences due to these mutations and elucidated the underlying molecular mechanisms using all-atom molecular dynamics simulations. We examine structural rearrangements of the RBD in the mutated variants leading to these differences. We also employ the ASGB method for a quantitative analysis of the binding mechanisms.

## Section 2. Materials and methods

### 2.1 Simulation set up

The three-dimensional structures of ACE2 and RBD domains of SARS-CoV-2 were obtained from the protein data bank PDB ID: 6M17^15^. We studied various mutated variants of this structure as listed in Table 1. The mutated structures were prepared using the CHIMERA software package.

**Table 1:** List of systems studied in this work.

| Mutation Variant | Mutations | System Size (number of atoms including water and ions) |
| --- | --- | --- |
| Wild Type | — | 194273 |
| Single Mutation | N501Y | 194280 |
| Double Mutation | E484Q, L452R | 232749 |
| Triple Mutation | N501Y, E484Q, L452R | 230658 |

Using the xLEaP module of AmberTools18^16^, the protein structures were enclosed in a water box with a buffer of 1.5 nm in all three directions. The systems were then charge neutralized by adding an appropriate number of Na^+^ ions. Additional Na^+^ ions and an equal number of Cl^−^ ions were further added to achieve physiological salt concentration of 150 mM. TIP3P water model^17^ was used for solvating the proteins. ff99sb-*ildn*^18^ force field was used to represent the inter- and intra-molecular interactions of the protein atoms. Interactions involving ions were described using the Joung Cheatham parameter set^19^.

### 2.2 MD simulation protocol

The solvated systems were initially energy minimized for 5000 steps using the steepest decent algorithm. The minimized structures were then heated from 0 to 300 K within a period of 50 ps with the solute atoms held fixed by a harmonic potential of strength 20 *kcal/mol/Å*^2^. After heating, the systems were subjected to a 2 ns long unrestrained equilibration. Finally, the equilibrated structures were subjected to 200 ns long production runs in the NVT ensemble. All the simulations were performed using the AMBER18 simulation package^16^. Periodic boundary condition was employed in all the three directions. SHAKE constraints were used for all bonds containing hydrogen allowing the use of a time step of 2 fs. Langevin thermostat with a collision frequency of 2 ps^*-*1^ was used for temperature regulation while isotropic Berendsen barostat with a coupling constant of 0.5 ps was used to regulate pressure in the NPT simulations. A cut-off of 10 Å was used to compute the short-range Lennard-Jones (LJ) interaction, while PME (Particle Mesh Ewald^20^) was employed for the calculation of long-range electrostatic interaction. The snapshots were visualized using VMD^21^ for analysis.

### 2.3 Binding energy calculation

The MMPBSA method^22,23^ employed in the MMPBSA.py module of AMBER18 was used to calculate the binding energies of the different ACE2-RBD complexes. The binding free energy difference (*ΔG*_*bind*_) of the SARS-CoV-2 RBD and ACE2 complex formation is calculated as^24^

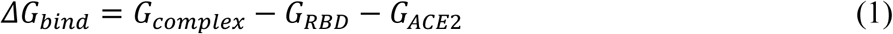

where *G*_*complex*_, *G*_*RBD*_, and *G*_*ACE2*_ represent the free energies of the RBD-ACE2 complex, individual RBD and individual ACE2 receptor, respectively. Eq. (1) can be decomposed into different interactions and can be written as

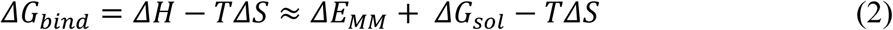

Here, *ΔE*_*MM*_ is the change in gas-phase molecular mechanics energy, *ΔS* is the entropy change (which has been neglected in this work), and *ΔG*_*sol*_ represents the solvation free energy change upon ligand binding. *ΔE*_*MM*_ can be further written as

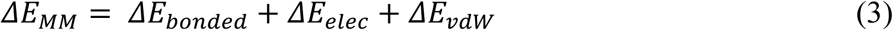

Here *ΔE*_*bonded*_, *ΔE*_*elec*_, and *ΔE*_*vdw*_ represent the change in bonded energies (bond, angle, and dihedral), electrostatic energies, and van der Waals energies upon ligand binding, respectively. *ΔG*_*sol*_ is the sum of the nonpolar solvation energy (*ΔG*_*SASA*_) and electrostatic solvation energy (*ΔG*_*PB*_), computed using the Poisson-Boltzmann method. *ΔG*_*SASA*_ can be written as,

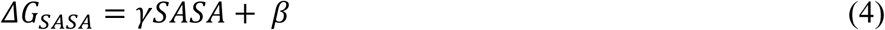

Here, *γ*(=0.00542 kcal/Å^2^) is the surface tension, while *β*=0.92 kcal/mol, and SASA represents the solvent-accessible surface area of the molecule.

### 2.4 Hotspot prediction by ASGB Method

The Alanine Scanning with MM/GBSA (ASGB) method is used on the last 10 ns of the 200 ns long MD simulation of each of the four RBD-ACE2 complex structures. In this method, all the residues lying within the 5 Å range of the protein–protein interaction interface are mutated. We use ASGB method with dielectric constants of 1, 3, and 5 for nonpolar, polar, and charged residues^24–26^, respectively. The binding free energy difference due to a point mutation was calculated by using the following relations,

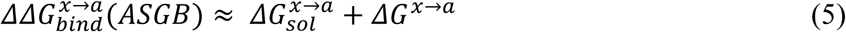

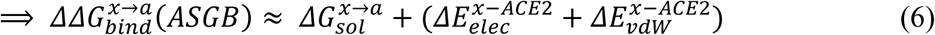

Here, 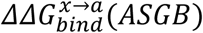 is the change in the binding free energy difference upon ligand binding due to a single point-mutation in RBD, while, 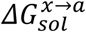 is the change in the solvation free energy difference upon ligand binding due to single point-mutation in RBD. 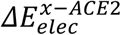 and 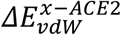 represent the difference in electrostatic and van der Waals energy between the mutated residue (*a*) in RBD with ACE2 to that of pre-mutated residue (*x*) in RBD with ACE2.

## Section 3. Results and Discussion

The structural stability of wild type, the single (N501Y), double (E484Q, L452R), and triple (N501Y, E484Q, L452R) mutants has been evaluated by analysing the RMSD fluctuations of backbone atoms and Cα atoms (Fig. 1). The single (N501Y) and double (E484Q, L452R) mutated variants have similar RMSD to wild type, while they show lower fluctuations than the triple (N501Y, E484Q, L452R) mutated variant. The RMSF fluctuations of all the 4 studied variants of SARS-CoV-2 RBD with ACE2 are shown in Fig. 2. The triple mutant (E484Q, L452R and N501Y) shows higher structural instability or fluctuations of RBD than the others.

**Fig. 1.**
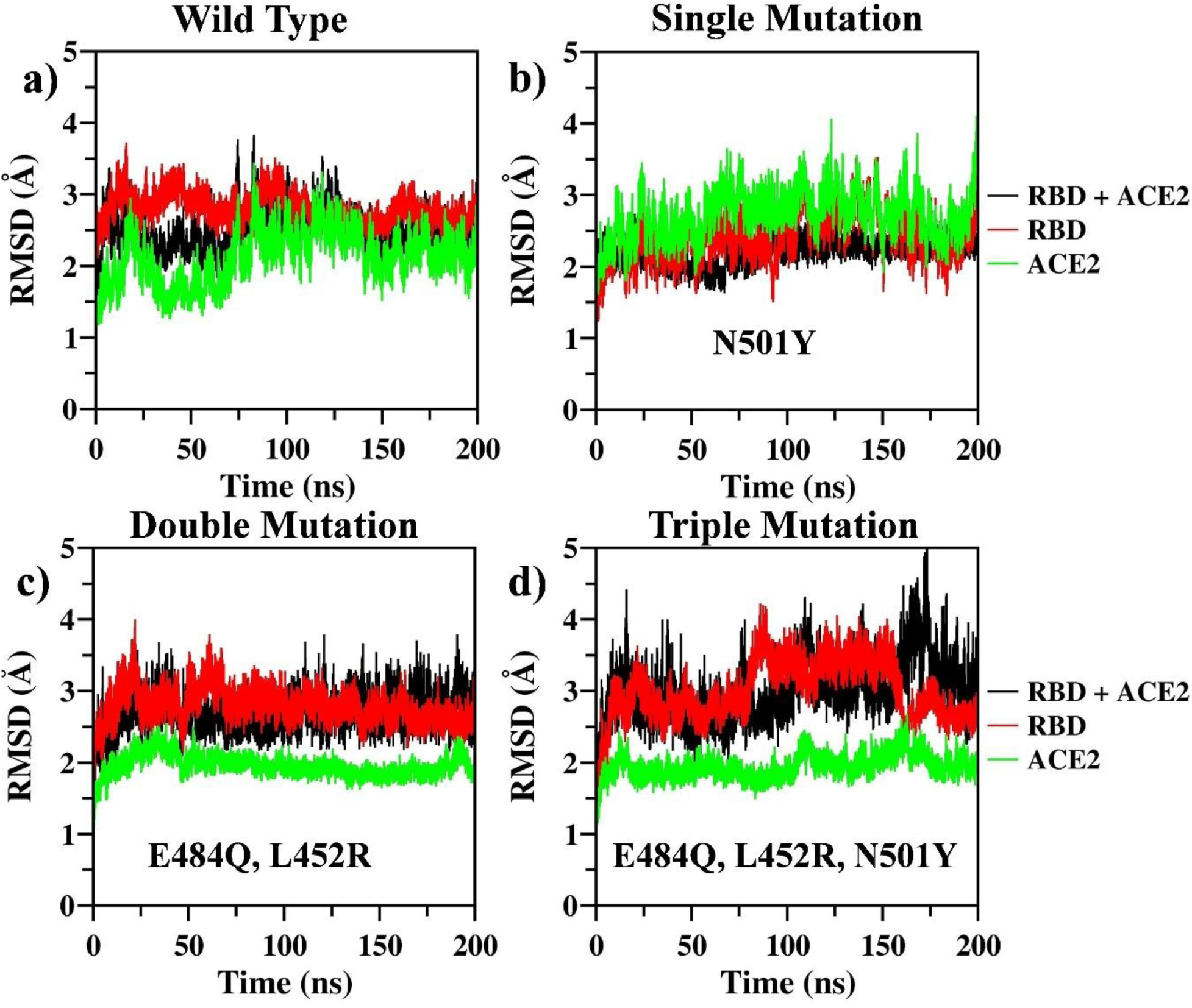
The RMSD values of the 200 ns long simulations of a) wild-type, b) N501Y single mutated, c) E484Q, L452R double mutated, and d) N501Y, E484Q, L452R triple mutated RBD variants, individually and in complex with ACE2. The RMSD value is small and similar for all the simulations, indicating the stability of the simulations performed.

**Fig. 2.**
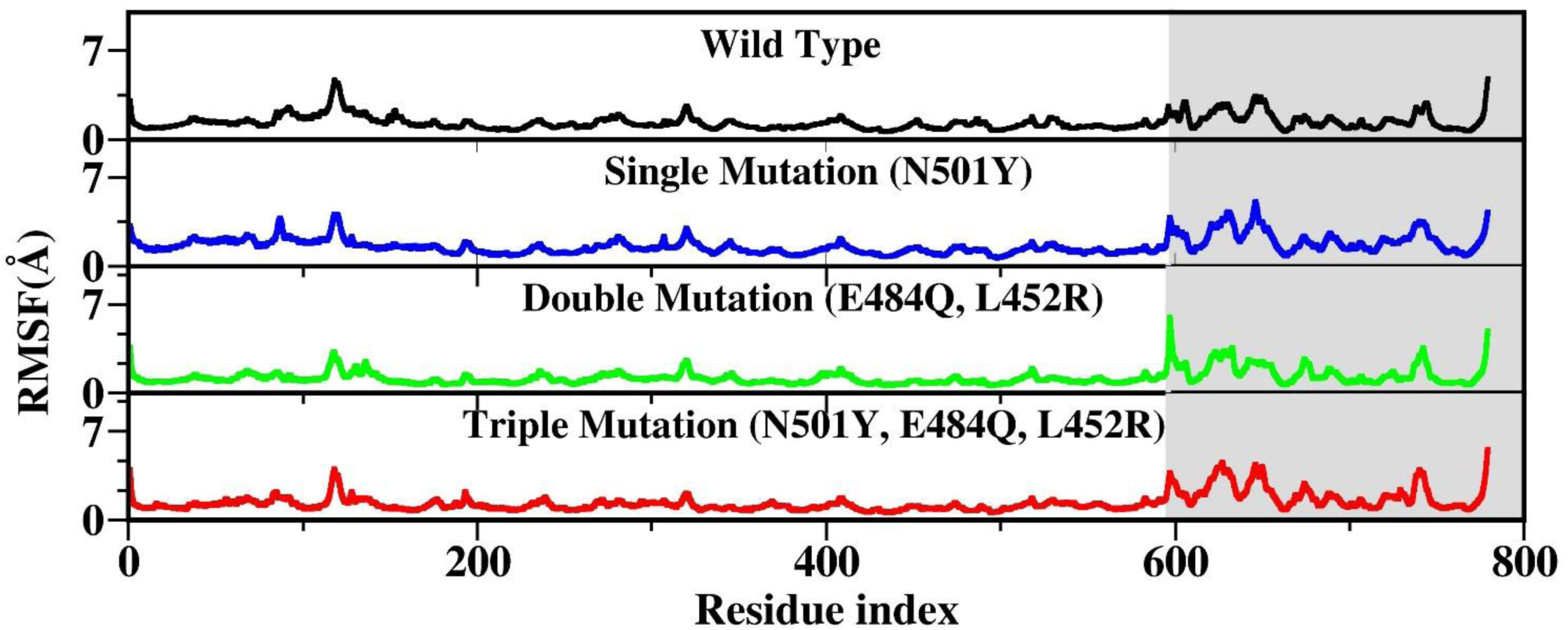
A comparison of residue-wise RMSF values for wild-type, N501Y single mutated, E484Q, L452R double mutated, and N501Y, E484Q, L452R triple mutated variants in simulations. The shaded area represents the residues of RBD while the unshaded area represents ACE2.

We present the superposed three-dimensional structures of the single (N501Y), double (E484Q, L452R), and triple (N501Y, E484Q, L452R) mutant variants with wild type SARS-CoV-2 RBD in Fig. 3 after 200 ns long MD simulations. The superposed 3-D structures indicate higher movements of the chains, including backbone atoms of the various mutated variants than that of the wild type RBD. The N501Y point mutation lies near the binding site to ACE2, while the other point mutation sites (E484Q and L452R) lie away from the binding sites. Fig. 4 shows the secondary structural changes of RBD for all the variants. The wild type RBD shows a transition from 25.1% sheet, 25.7% turn, and 49.2% coil to 3.3% helix, 25.8% sheet, 22.5% turn, and 48.4% coil during 200 ns simulation. The introduction of N501Y mutation has changed the final secondary structure content to 2.7% helix, 30.6% sheet, 24.0% turn, 40.4% coil, and 2.2% 3-10 helix. In comparison, the double (E484Q, L452R) and triple (N501Y, E484Q, L452R) mutants show 24.7% sheet, 22.5% turn, 50.5% coil, and 2.2% 3-10 helix, and 3.3% helix, 24.2% sheet, 23.6% turn, and 48.9% coil after 200 ns, respectively. There are more transitions in the secondary structure for the triple (N501Y, E484Q, L452R) mutant variant than the double mutant (E484Q, L452R) and the wild-type. This indicates the structural rearrangements induced by these mutations even while not in direct contact with ACE2. The double (E484Q, L452R) and triple (N501Y, E484Q, L452R) mutants generate large conformational switches that orient the RBD towards the N-terminal of ACE2 and α-helix region. This conformational rearrangement is found to generate more contacts with ACE2.

**Fig. 3.**
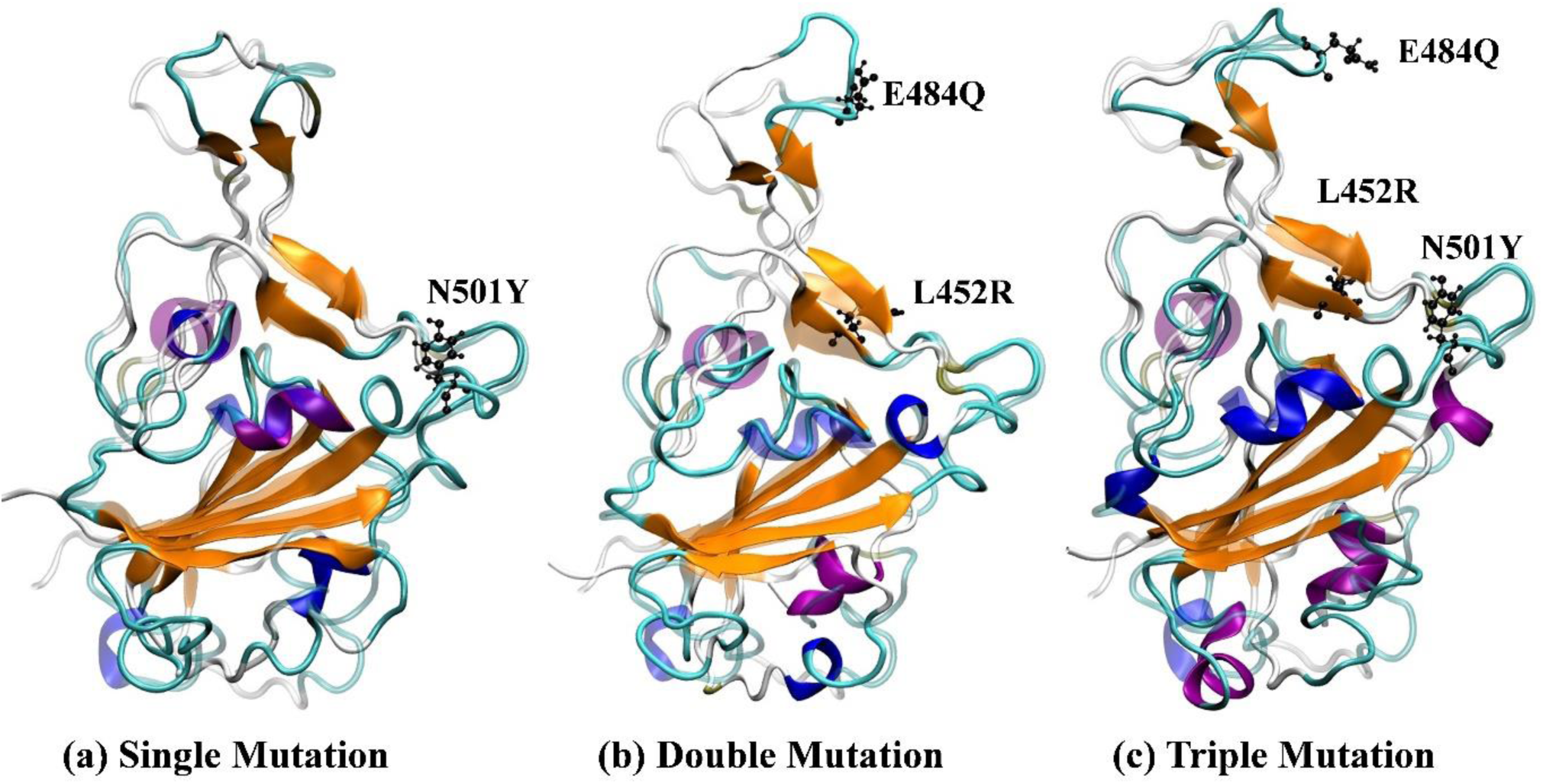
Superimposed structures of wild-type (transparent representation) and a) N501Y single mutated, b) E484Q, L452R double mutated, and c) N501Y, E484Q, L452R triple mutated (solid representation) RBD structures. The mutated residues are shown in black in CPK representation. The mutated structures contain many secondary structure deviations from the wild type.

**Fig. 4.**
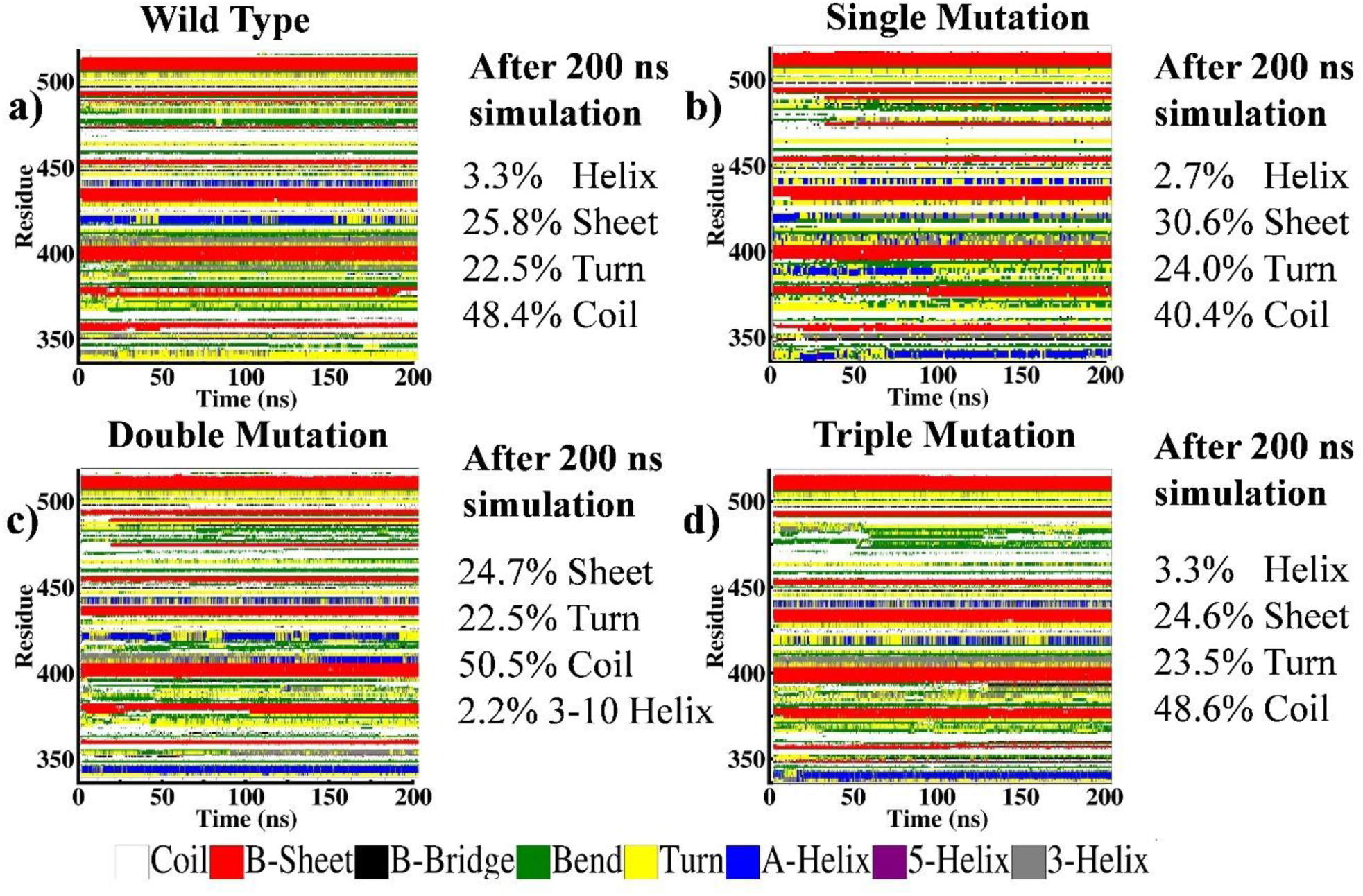
The secondary structure transition of a) wild-type, b) N501Y single mutated, c) E484Q, L452R double mutated, and d) N501Y, E484Q, L452R triple mutated SARS-CoV-2 RBD after 200 ns simulations.

We have calculated the number of hydrogen bonds between RBD-ACE2 for wild type and mutated variants (Fig. 5). A hydrogen bond (H-bond) is defined to exist between a hydrogen atom involved in a polar bond and an electronegative acceptor (A) using a distance cut-off of 3.5 Å and a D—H…A angle cut-off of 35°, D being the donor. We have calculated the number of active H-bonds over the last 100 ns of the 200 ns long trajectories. Clearly, there are more hydrogen bonds formed between the RBD and ACE2 for double (E484Q, L452R), and triple (N501Y, E484Q, L452R) mutants than the wild type variant. However, the single N501Y mutation shows the least number of intact H-bonds over the last 100 ns of the 200 ns long MD simulations. This indicates the weaker binding of N501Y variant than double (E484Q, L452R) and triple (N501Y, E484Q, L452R) mutant variants.

**Fig. 5.**
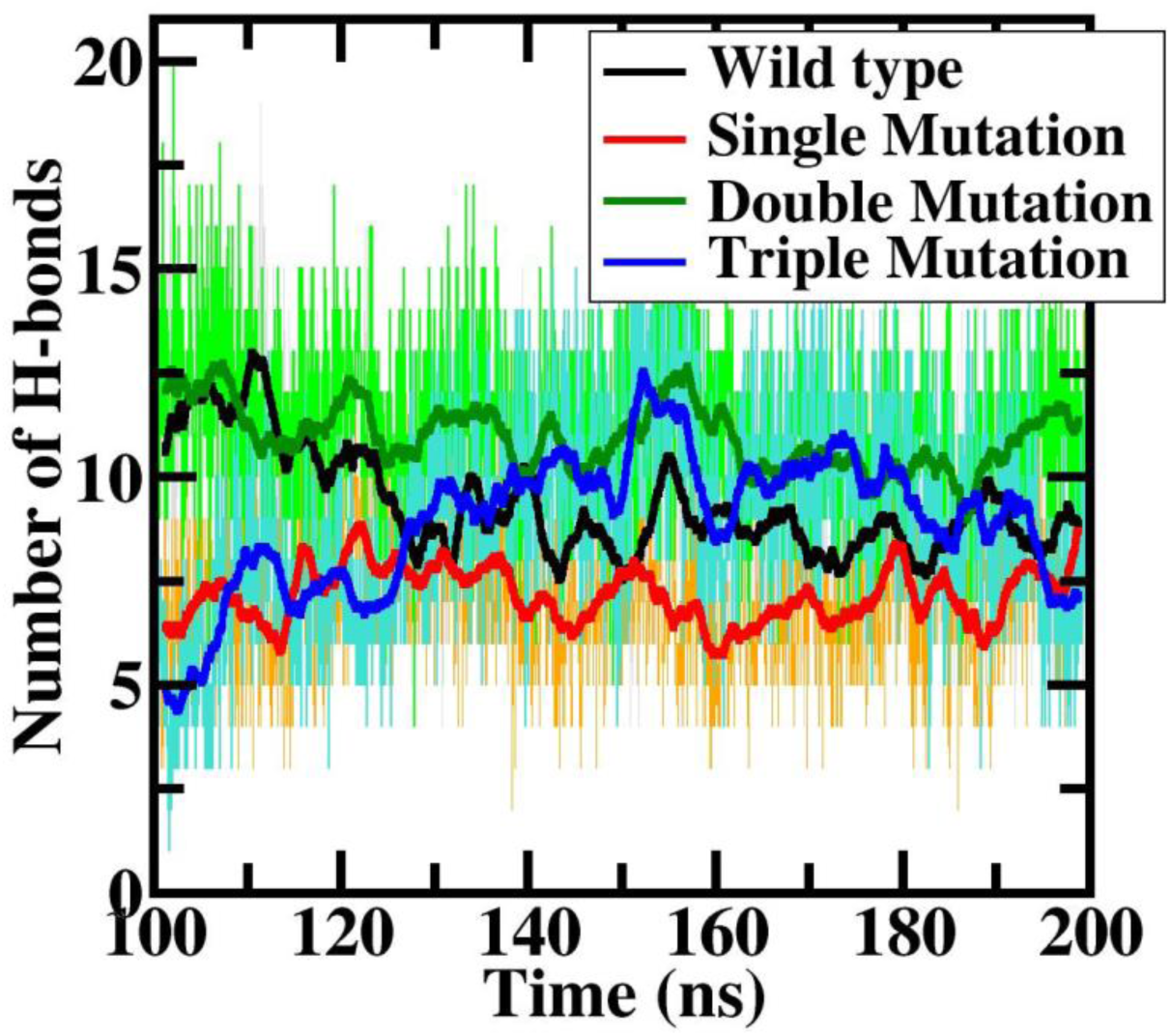
A comparison of the number of H-bonds between the ACE2 and RBD of wild-type, N501Y single mutated, E484Q, L452R double mutated, and N501Y, E484Q, L452R triple mutated SARS-CoV-2 during the last 100 ns of 200 ns long MD simulations.

Using MM-PBSA method, we compute the total binding energy of RBD with ACE2 (Fig. 6). The binding energy calculated using the MMPBSA method over the last 100 ns of the 200 ns long MD simulations shows a higher affinity for double (E484Q, L452R) mutant (−98.6 +/- 7.3 kcal/mol) than wild type (−59.7 +/- 9.6 kcal/mol), single (N501Y) mutant (−58.0 +/- 8.0 kcal/mol) and triple (N501Y, E484Q, L452R) mutant (−79.8 +/- 9.2 kcal/mol). If this increased binding were to translate to increased infectivity, it may explain the high infection rate of the B.1.617 variant which recently caused the second wave of COVID infections in India.

**Fig. 6.**
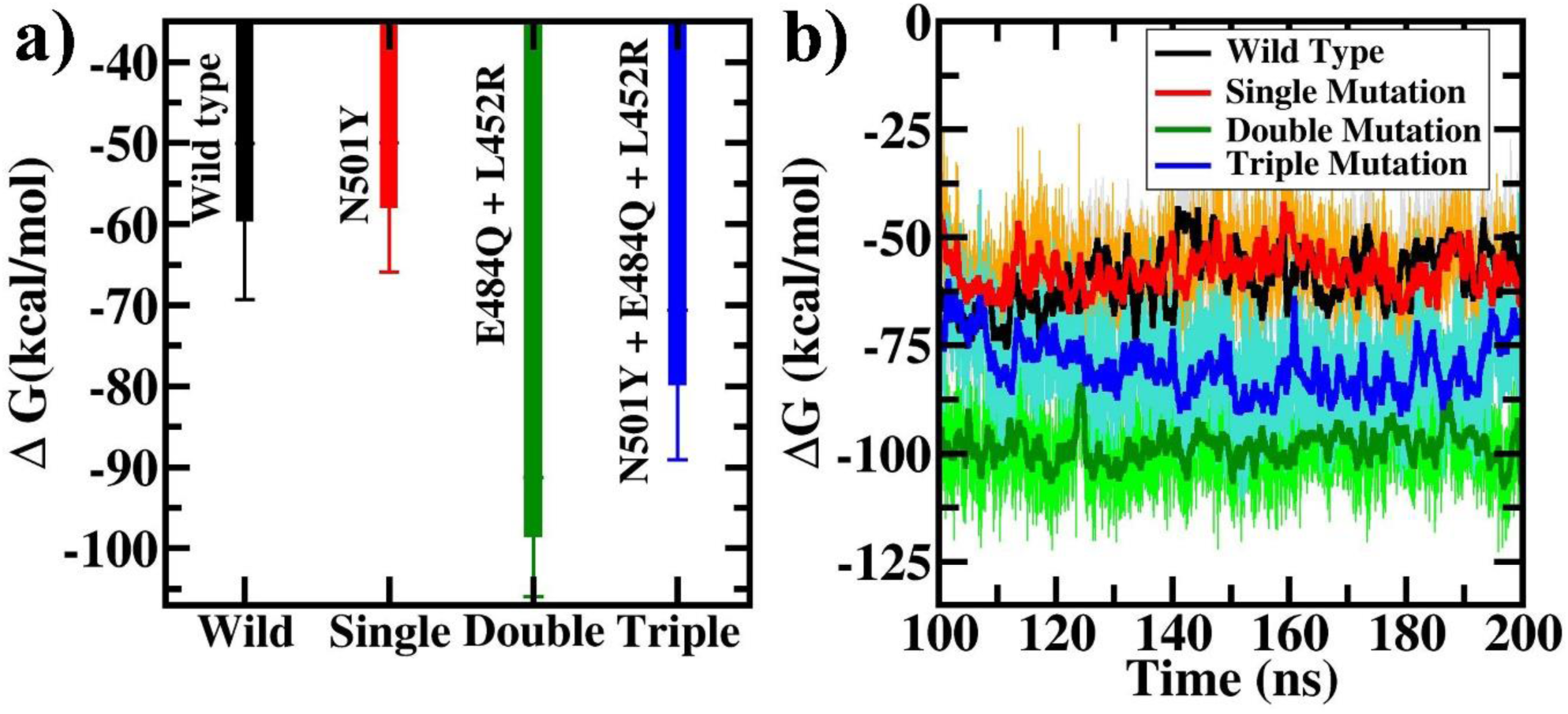
(a) The binding energies of ACE2 with wild-type compared to that with N501Y single mutated, E484Q, L452R double mutated, and N501Y, E484Q, L452R triple mutated RBD computed using MMPBSA simulations. (b) The variation of binding energy during the last 100 ns of the corresponding 200 ns long simulations.

To gain further insight into the mechanism causing these changes in the binding affinities, we have used the alanine scanning method (ASGB) to predict the hotspot residues in RBD involved in ACE2-RBD binding. The residues Asn709, Tyr710, Ile733, Val744, Glu745, Phe751 and Asn762 show more than 2 kcal/mol energy differences. The reason for high binding energy from these residues is their positioning at the ACE2 binding site. Among these residues, we found that Glu745 contributed more due to its hydrogen bond formation with the Phe10 of ACE2. Further, the contribution from residues 669R, 678K, 710Y, 747F, 754Q, 759Q, 761T and 766Y are higher for the wild type ACE2-RBD (Fig. 7). Large conformational changes are caused due to the mutations in RBD that increase the number of residues that interact with ACE2 receptor. We find that 747F is a key contributor to the binding energy of all the mutated variants of RBD we considered. The total contribution from the hotspot residues is -20.84 kcal/mol and single and double mutant were identified as -29.94 kcal/mol and -36.1 kcal/mol. The triple (N501Y, E484Q, L452R) mutant variant shows a higher binding affinity than wild type (−33.06 kcal/mol). As shown in Table 2, the contribution from each residue is found to be higher for the mutated variants than the wild type. Moreover, the double mutated (E484Q, L452R) variant shows the highest binding affinity. We note that ASGB does not consider the repulsive interactions which may lead to a difference between the computed binding energies using MMPBSA and ASGB. Nevertheless, the qualitative results obtained using ASGB agree well with MMPBSA and indicate that the double mutant (E484Q, L452R) RBD spike has the highest binding affinity for ACE2 among the variants considered.

**Fig. 7.**
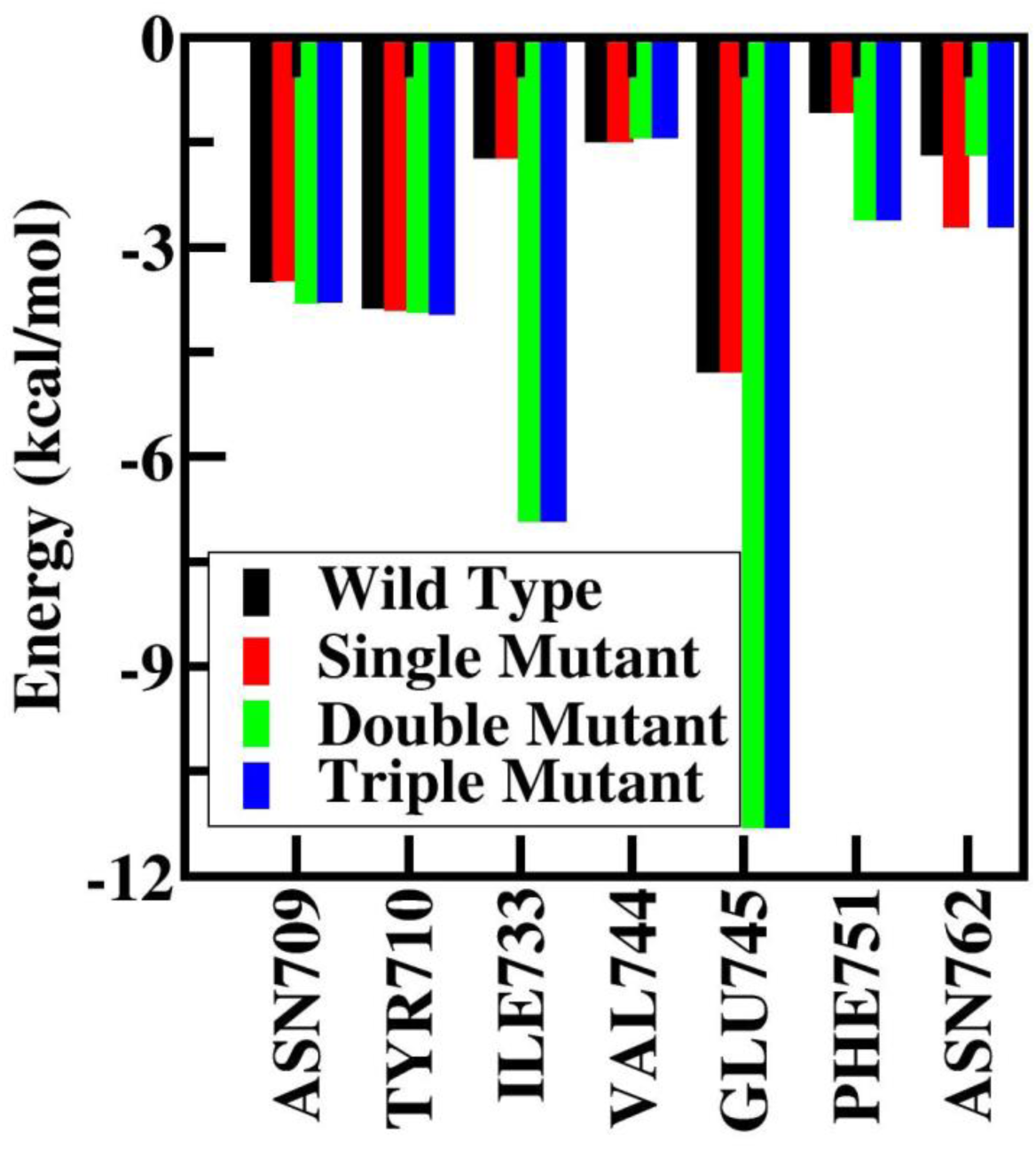
Hotspot residues of ACE2 binding to wild type, N501Y single mutated, E484Q, L452R double mutated, and N501Y, E484Q, L452R triple mutated RBD obtained using the ASGB calculation.

**Table 2.**
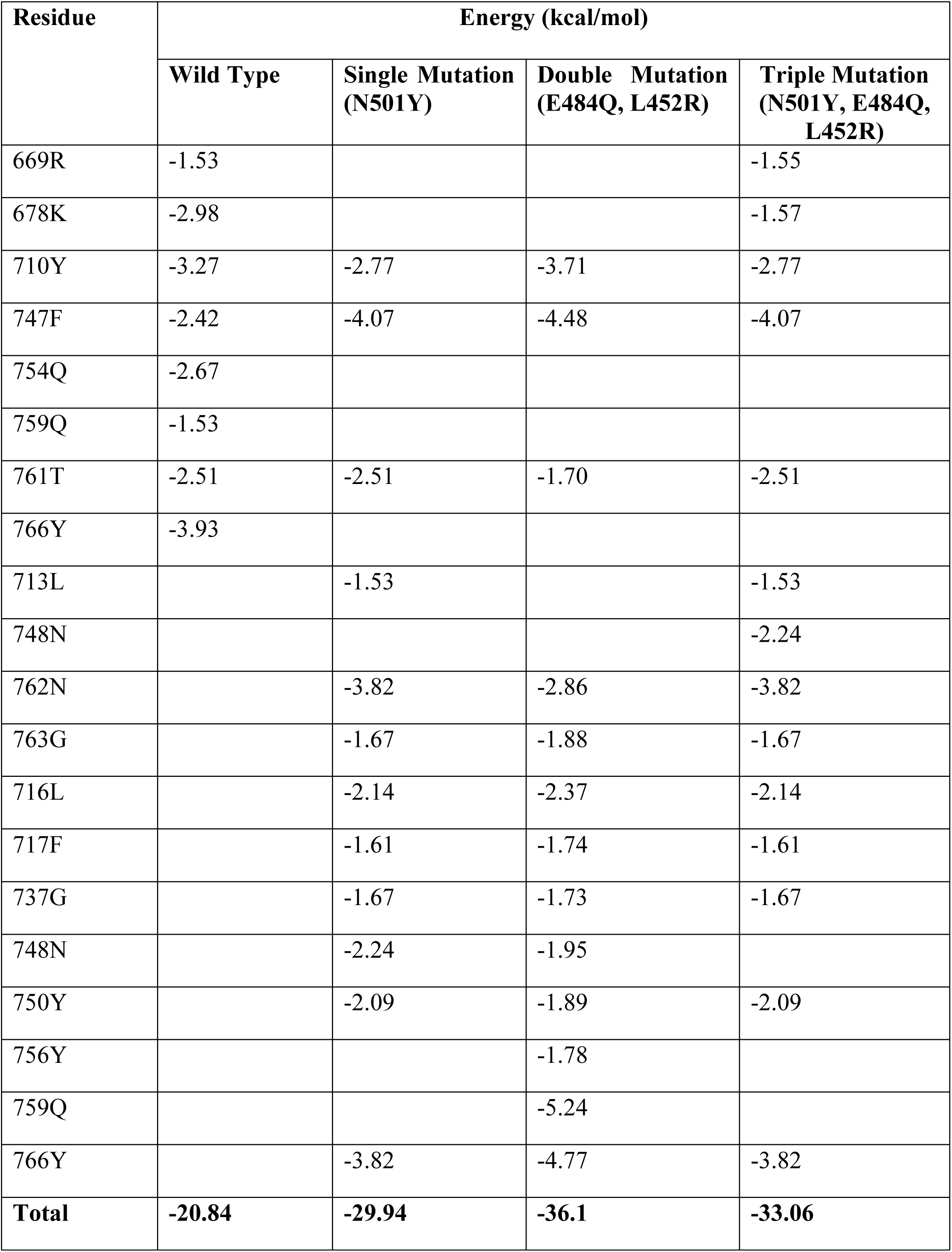
Calculated binding free energy from key residues of RBD domain binding to ACE2. Energy values are in kcal/mol and energy values above -1.5 kcal/mol are considered.

Next, we have compared our estimated binding energy data with available experimental values. Experimental binding affinity values of mutated SARS-CoV-2 RBD with ACE2 are quite sparse. There are two major experiments—one by Laffeber et al.^27^ and the other by Kim et al.^28^—that reported the equilibrium dissociation constant (*K*_*D*_) for several SARS-CoV-2 mutants. Binding energy is then calculated from *K*_*D*_ using the relation, Δ*G* = −*RT In K*_*D*_. As shown in table 3, the difference in binding energy (ΔΔ*G*) of all SARS-CoV-2 mutant variants is higher than the wild-type, which agrees well with the experimental trend, although, the simulation results overestimated the experimental data. Such overestimation of the binding energy is quite usual in simulations and is often related to the different force-fields used and the overestimation of the van der Waals interaction energy^26^.

**Table 3.**
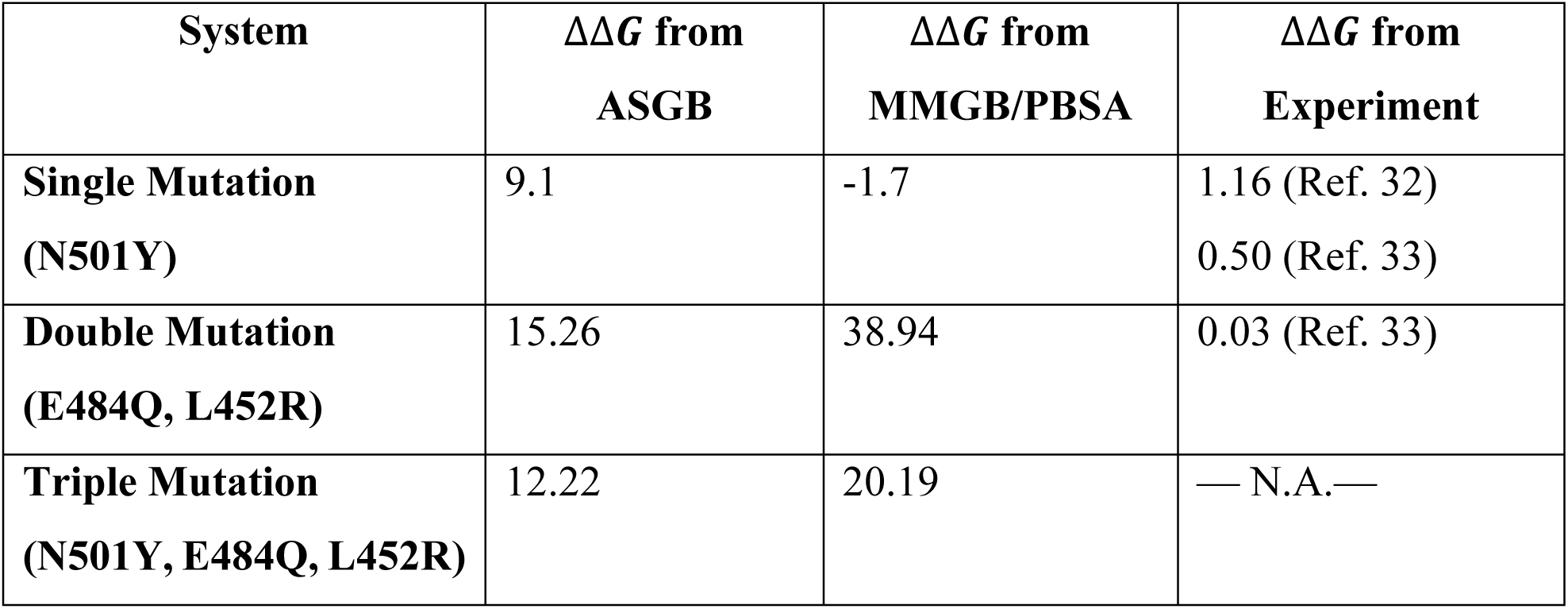
Calculated differences in binding free (ΔΔ*G*) energies of mutated SARS-CoV-2 variants and wild-type RBD and comparison with experiments. ΔΔ*G* of different SARS-CoV-2 mutated variants is calculated with respect to the wild type.

## Section 4. Conclusion

We have performed extensive MD simulations of the binding of wild-type SARS-CoV-2 RBD, and its single (N501Y), double (E484Q, L452R) and triple (N501Y, E484Q, L452R) mutated variants with human ACE2. We have used advanced free energy methods to calculate the associated binding energies. The double mutant (E484Q, L452R) variant shows the highest affinity followed by the triple mutant (N501Y, E484Q, L452R), the wild-type, and the single (N501Y) mutant. The higher binding affinity arises due to the structural changes induced by the mutations. Using the alanine scanning method, the residues of the RBD contributing the most to the binding energy difference were identified. The residue 747F was identified as the key contributor in all the mutated variants we studied. For the double (E484Q, L452R) mutated variant, the 759Q and 747F residues contribute the most, while for the triple (N501Y, E484Q, L452R) mutated variant, the residues 747F, 762N and 710Y contribute the most. This study provides new mechanistic insights into the interactions between the wild-type and various mutated variants of SARS-CoV-2 with the ACE2 receptor, enhancing our understanding of how the variants may acquire increased transmissibility. The structural insights gained may also help design site-specific therapeutic molecules that may work against multiple variants.

## Acknowledgements

We thank DST, India for computational support through TUE-CMS computational facility. A.A. thanks MHRD, India for the generous fellowship. S.N thanks CSIR, India for the generous fellowship. N.M. thanks Indian Institute of Science (IISc) for support through IoE (Institute of Eminence) postdoctoral scheme.

